# Engineering Functional Pseudo-Islets of Defined Sizes from Primary Murine Cells Using PEG Microwell Devices

**DOI:** 10.1101/2020.02.07.939538

**Authors:** Kelly MT Shekiro, Thomas H Hraha, Abigail B Bernard, Richard KP Benninger, Kristi S Anseth

**Author notes:** Corresponding author, Full postal: Department of Chemical and Biological Engineering, University of Colorado, UCB 596, Boulder, CO 80309.

## Abstract

A major limitation of islet transplantation as a therapy for treating Type 1 Diabetes is eventual graft failure, which can be partially attributed to islet cell death. When cultured *in vitro,* cells in the center of large islets show increased necrosis and exhibit decreased viability and insulin secretion compared to smaller islets. Given the necessity of β-cell-to-β-cell coupling for the physiological response to glucose, a technique to re-aggregate primary islet cells or cells derived from progenitor cells into small clusters of defined sizes may prove advantageous for promoting function upon transplantation. Here, hydrogel microwell arrays were utilized to generate 3-dimensional pseudo-islets from primary murine islets. Pseudo-islets ranged from 50 to 100 μm in diameter as controlled through the microwell dimensions, and contained β-, α-, and δ-cells with ratios similar to those in whole murine islets. Over two weeks in culture, pseudo-islets remained highly viable and responsive to glucose. Intracellular calcium flux showed more robust and coordinated dynamics at high glucose and decreased activity at low glucose compared to age-matched wild-type islets. Therefore, microwell devices can control the aggregation of cells isolated from primary islets to produce islet-like clusters that are functionally similar to freshly isolated islets, and may provide a technique to create improved cellular therapies for Type 1 Diabetes.

## 1. Introduction

Islet transplantation is a promising therapy for the treatment of type 1 diabetes that may eliminate patients’ need for frequent blood glucose monitoring and exogenous insulin therapy. Indeed, transplantation via the Edmonton protocol has offered improved glycemic control and reduced insulin dependence post-transplantation [1,2]. Despite a specialized immunosuppressive regime to modulate the immune response to the transplant [1], many patients require a second or third transplantation, and eventually experience graft failure and must resume insulin dosing [2,3]. While the immune response likely affects graft function, lack of vasculature at the transplant site is also a major contributing factor to graft failure [4].

Without the direct delivery method provided by the islet microvasculature, oxygen and other nutrients must diffuse through the outer, dense cell mass to reach the inner-most cells [5]. The impact of diffusion-limited nutrient transport can be observed by the increase in cell necrosis at the center of larger islets, typically >125 μm diameter, which is not observed in smaller islets [5,6]. In addition, smaller islets display increased viability and functional response [7,8], suggesting that islet size is a critical factor in the development of an effective transplantation-based therapy for type 1 diabetes. Given the limited supply of high quality donor tissue [9,10] and that alternative cell sources, such as β-cells derived from stem cells [11–13], have not yet been proven to work on a large scale, methods to improve the efficiency of β-cell transplantation are needed.

To improve accessibility of nutrients and reduce diffusion barriers, researchers have proposed the use of single β-cells within a cell delivery scaffold [14]. However, the necessity of cell-cell contact between β-cells to maintain a normal response to glucose, whether through direct cell contact [15–19] or through cell mimicry [20,21], is well documented. Insulin secretion is a highly dynamic process where the metabolism of glucose within the β-cell leads to changes in cell membrane polarization, allowing for an influx of calcium which triggers insulin secretion [22]. During these depolarization events, insulin release between neighboring cells is synchronized through gap junction coupling [23–25]. Synchronization between cells at high glucose results in bursts of insulin [26] into the blood plasma, which better controls blood glucose levels [27] and the suppression of insulin release at low glucose when it is not needed [28,29].

Given the importance of β-cell-to-β-cell contact, several methods have been developed towards the goal of creating β-cell aggregates. The most common and easiest method is spontaneous aggregation of β-cells either in static or orbital culture [14,19,30–32]; however, this technique affords very little control over the final aggregate size and shape. Another method uses hanging drops, where single β-cells are placed in drops on a surface and then inverted and cultured, leading to aggregation of the cells within the drop [33]. This protocol produces aggregates of defined size, but is limited by the inability to produce aggregates across a range of desired sizes and is fairly labor intensive to execute. Microcontact printing of proteins has been used to create islands of multi-cellular β-cell clusters on glass surfaces [34,35]. While the dimensions of the protein island dictate final aggregate size and allows for a range of sizes, these clusters are either monolayers of cells or a few cell layers high at most, and removal of these β-cell clusters from the surface for 3D studies or transplantation has not yet been demonstrated. More recently, microwell scaffolds made from glass micromolds [36], poly(ethylene oxide terephthalate) and poly(butylenes terephthalate) copolymer thin films and electrospun meshes [37], and poly(ethylene glycol) (PEG) microwell arrays [38] have been used to create β-cell aggregates of defined sizes. These techniques allow for the facile creation of a large number of aggregates across a range of sizes dictated by the microwell scaffold geometry used.

In this work, we demonstrate the use of PEG microwell arrays to create three-dimensional pseudo-islets from primary murine islets. Microwell device size was varied to create pseudo-islets of multiple diameters. Immunofluorescence staining of three hormones found in different islet cells was performed to investigate the cellular composition of pseudo-islets. Whole and pseudo-islets were maintained in *in vitro* culture over two weeks post-isolation to assay for viability and evaluate static insulin secretion as a measure of pseudo-islet functionality. Additionally, the dynamics of insulin secretion in pseudo-islets was studied to understand the extent of β-cell coupling within pseudo-islets.

## 2. Materials and Methods

### 2.1 Materials

All chemical reagents were purchased from Sigma-Aldrich, unless otherwise noted. All cell culture reagents were obtained from Life Technologies.

### 2.2 Microwell Device Fabrication

Poly(ethylene glycol) (PEG) hydrogel microwell arrays were fabricated through contact photolithography as previously described [38]. A pre-polymer solution contained 10.8 wt% PEG diacrylate (PEGDA, M_n_ ∼3,000 Da, synthesized from hydroxyl-terminated PEG as described [39], 4.2 wt% poly(ethylene glycol) monoacrylate (PEGA, M_n_ ∼400Da; Monomer-Polymer and Dajac Labs), 0.05 wt% of the photoinitiator 4-(2-hydroxyethoxy)phenyl-(1-hydroxy-2-propyl)ketone (Irgacure 2959; Ciba) and Dulbecco’s phosphate buffered saline (PBS). Piranha cleaned glass slides were treated with (3-acryloxypropyl)trimethoxysilane (Gelest) by chemical vapor deposition [40] to create acrylated surfaces. The pre-polymer solution was then placed between the acrylated glass surface, sized to fit into a 48-well plate, and a chrome photomask (Photo Sciences, Inc.) and irradiated with collimated, broadband UV light (λ = 350-500 nm, I ∼ 70 mW/cm^2^; Oriel Instruments) for 60 sec to allow for polymerization in the regions not shaded by the photomask. The photomask patterns created square well arrays, either 100 μm (w100) or 200 μm (w200) wide and 100 μm tall, controlled by adjusting the distance between the glass slide and photomask. The devices were rinsed in water to remove any unreacted components and sterilized by dipping in 70% ethanol and at least 2 hours of exposure to a germicidal UV light prior to use with cells.

### 2.3 Pseudo-islet Formation

Female, retired breeder Balb/cByJ mice (Jackson Laboratories) were housed in a centralized animal care facility at the University of Colorado Anschutz Medical Campus. NIH guidelines for the care and use of laboratory animals (NIH Publication #85-23 Rev. 1985) were followed and all experiments involving mice were approved by the University of Colorado Institutional Animal Care and Use Committee. Islets were isolated at the Diabetes and Endocrinology Research Center at the Barbara Davis Center for Childhood Diabetes in Aurora, CO and transported to Boulder, CO on ice. Islets were cultured overnight in untreated culture dishes in RPMI media with 20% fetal bovine serum, 1% penicillin-streptomycin and 0.2% Fungizone. A single cell suspension of the primary cells was obtained by sequential 0.01% trypsin-EDTA digestion. Single cells were suspended at a density of 400,000 or 1,000,000 cells/mL for w100 and w200 devices, respectively, and 500 μL of the cell suspension was seeded into microwell arrays in 48-well cell culture plates. The cells and devices were centrifuged at 160 rcf for 2.5 min; unseeded cells were re-suspended and the centrifugation was repeated before 2 hours of orbital culture at 35 rpm in an incubator (37°C, 5% CO_2_) followed by 5 days of static culture. Pseudo-islets were removed by rinsing the devices with media and then placing devices vertically into a conical tube and centrifuging at 200 rcf for 5 min to assist with removal.

### 2.4 Pseudo-islet Size Determination

To determine the effect of device dimension on pseudo-islet size, brightfield images were collected of pseudo-islets after removal from the devices on a Nikon Eclipse TE2000-S microscope (Nikon). ImageJ (NIH) software was used to fit an ellipse to the pseudo-islet and the major axis was measured.

### 2.5 Live/Dead

Viability of whole and pseudo-islets was observed by staining with a Live/Dead Viability/Cytotoxicity kit (Life Technologies). Briefly, cells were incubated in phenol-red free RPMI with 2 μM calcein-AM and 4 μm ethidium homodimer-1 for 45 min prior to imaging via confocal microscopy (Zeiss NLO LSM 710, Carl Zeiss) where live cells stain green and dead cells are indicated by red nuclear staining.

### 2.6 Immunostaining

The cellular composition of whole and pseudo-islets was investigated by staining the islets for insulin, glucagon and somatostatin. Whole and pseudo-islets were fixed in 4% paraformaldehyde (PFA) for 25 min at room temperature prior to permeabilization in 0.3% Triton-X 100 in PBS for 3 hours. After blocking overnight at 4°C in 5% bovine serum albumin (BSA)/0.15% Triton-X 100 in PBS, the samples were equilibrated in antibody buffer. The antibody buffer contained 1% BSA/0.2% Triton-X 100 in PBS. Samples were incubated in primary antibodies overnight at 4°C, washed in antibody buffer, incubated in secondary antibodies overnight at 4°C and washed again. Antibodies used include guinea pig anti-insulin (Abcam ab7842, 1:50), mouse anti-glucagon (Abcam ab10988, 1:100), rabbit anti-somatostatin (Santa Cruz, sc-13099), Alexa Fluor 633 goat anti-guinea pig (Life Technologies A21105, 10 μg/mL), Alexa Fluor 488 donkey anti-mouse (Life Technologies A21202, 10 μg/mL), and Alexa Fluor 555 goat anti-rabbit (Life Technologies A21430, 10 μg/mL). 4’-6-diamidino-1-phenylindole (DAPI, Life Technologies) was applied at 1 μg/mL for 90 minutes to stain the nuclei and the samples were washed and mounted with AquaPoly/Mount (Polysciences). Whole and pseudo-islets were imaged on a Zeiss NLO LSM 710 laser scanning confocal microscope (Carl Zeiss).

Immunostaining was also completed to look for connexin-36, which forms gap junctions between β-cells. Whole and pseudo-islets were entrapped within a hydrogel (10 wt% PEGDA, M_n_ ∼10 kDa, and 1.7 mM LAP, polymerized with 365 nm light at ∼ 5 mW/cm^2^ for 3 min) and incubated in media overnight. The samples were fixed in 4% PFA for 25 min at room temperature. The gels were soaked in HistoPrep (Fisher Scientific) overnight and then directly frozen in liquid nitrogen and stored at -20°C. Samples were cryosectioned into 50 μm slices and mounted onto ColorFrost Plus slides (Fisher Scientific). The sections were blocked with 5% BSA in PBS prior to incubation overnight at 4°C with primary antibody (rabbit anti-connexin-36, Life Technologies 36-4600, 1:50) in 1% BSA. After washing, the secondary antibody (Alexa Fluor 488 goat anti-rabbit, Life Technologies A11008, 10 μg/mL) was applied overnight at 4°C. DAPI was applied at 1 μg/mL for 15 minutes to stain the nuclei and the samples were washed and imaged on a Zeiss NLO 710 laser scanning confocal microscope (Carl Zeiss).

### 2.7 Insulin Secretion

Static insulin secretion assays were completed to test the functional response of the pseudo-islets. Whole and pseudo-islets were conditioned in Krebs-Ringer Buffer (KRB, 128.8 mM NaCl, 5 mM Na HCO_3_, 1.2 mM KCl, 2.5 mM KH_2_PO_4_, 1.2 mM MgSO_4_, 10 mM Hepes, 0.1% bovine serum albumin, pH = 7.4) with 2 mM glucose for 1 hour before incubation in either 2 mM glucose (low glucose) or 20 mM (high glucose) KRB for an additional hour. The supernatant was collected and the islets were lysed with 1% Triton-X 100. The insulin content of samples and the lysate was analyzed using a sandwich ELISA per the manufacturer’s protocol (Mercodia).

### 2.8 Dynamic Calcium Microscopy

β-cell calcium dynamics were observed by staining whole and pseudo-islets with 4 μM Fluo4-AM (Life Technologies) in imaging buffer (125 mM NaCl, 5.7 mM KCl, 2.5 mM CaCl_2_, 1.2 mM MgCl_2_, 10 mM Hepes, 0.1% bovine serum albumin, pH 7.4) for 60-90 min at room temperature on an orbital shaker. Samples were loaded into a polydimethylsiloxane (PDMS) microfluidic device (fabricated as described [41]), and exposed to imaging buffer containing 2 mM or 11 mM glucose, allowed to equilibrate for 7 min and imaged in one second intervals. A Nikon Eclipse Ti microscope with a humidified environmental chamber at 37°C was used for real-time imaging.

### 2.9 Quantitative Image Analysis

MATLAB (MathWorks) was used to analyze real-time calcium fluorescent images using methods that have been previously detailed [42]. To quantify the robustness of the calcium oscillations, the Fourier transform of the fluorescent intensity time-courses was taken, and the peak AC component was normalized to the DC (0 frequency) component. The area of synchronization was calculated by computing a cross-correlation coefficient between the average fluorescence intensity of the entire islet (as reference) and each pixel of the image. The maximum cross-correlation value (equivalent to the degree of overlap, where 1 indicates perfect correlation and zero indicates no correlation) was determined for each pixel, and correlations greater than an arbitrary threshold of 0.75 were considered synchronized. The fractional area of the islet with synchronized behavior was then calculated. In order to quantify the level of activity within an islet at low glucose, a reference ‘silent’ cell was selected and a pixel was defined as active if the standard deviation of its calcium trace was greater than three standard deviations from the reference cell’s. The fractional area of the islet defined to be active was then calculated.

### 2.10 Data Analysis

All results are from at least two independent isolations of 8 mice each. A minimum of two replicates per isolation were tested for each condition depending on the number of whole or pseudo-islets available. For insulin secretion studies, 10 whole islets or pseudo-islets were pooled for a single data point. Pseudo-islet diameter is reported as mean ± standard deviation due to the large number of measurements taken; all other data are presented as mean ± standard error. Data was analyzed with Graphpad Prism 5 software, using a t-test or ANOVA with a Bonferroni correction for multiple comparisons. Statistical significance was noted when p < 0.05.

## 3. Results

### 3.1 Creation of three-dimensional pseudo-islets of defined sizes from primary murine islets

Previous work using hydrogel microwell arrays to aggregate MIN6 β-cells has demonstrated their utility in creating a large number of consistent multi-cellular aggregates across a range of user-defined sizes [38]. Here, contact photolithography was used as previously described [38] to fabricate PEG based devices that are robust and easy to handle during pseudo-islet formation. The microwell device pre-polymer solution was formulated to polymerize quickly and minimize volume changes after polymerization allowing for high fidelity of pattern transfer from the chrome mask to the microwells. For this work, square microwells were 100 μm deep and either 100 μm (w100) or 200 μm (w200) wide. Approximately 1225 wells fit on each w100 device and w200 devices had ∼550 wells.

After isolation from mice and overnight suspension culture, whole islets were dissociated into single cells prior to seeding into microwells (Figure 1). A seeding density of 400,000 or 1,000,000 single primary cells/mL was found to amply fill the two sizes of microwell devices tested here (Figure 2A). Over the course of five days in culture, the cells transitioned from distinct single cells into multicellular clusters that remained intact after removal from the device and placed in suspension culture (Figure 2A). The pseudo-islets, as well as control whole islets, were maintained in suspension culture for analysis at day 7, one week after isolation, and day 14, two weeks after isolation. Given that the pseudo-islets take 6 days to aggregate within the microwell devices, the earliest they could be assayed was on day 7.

**Figure 1.**
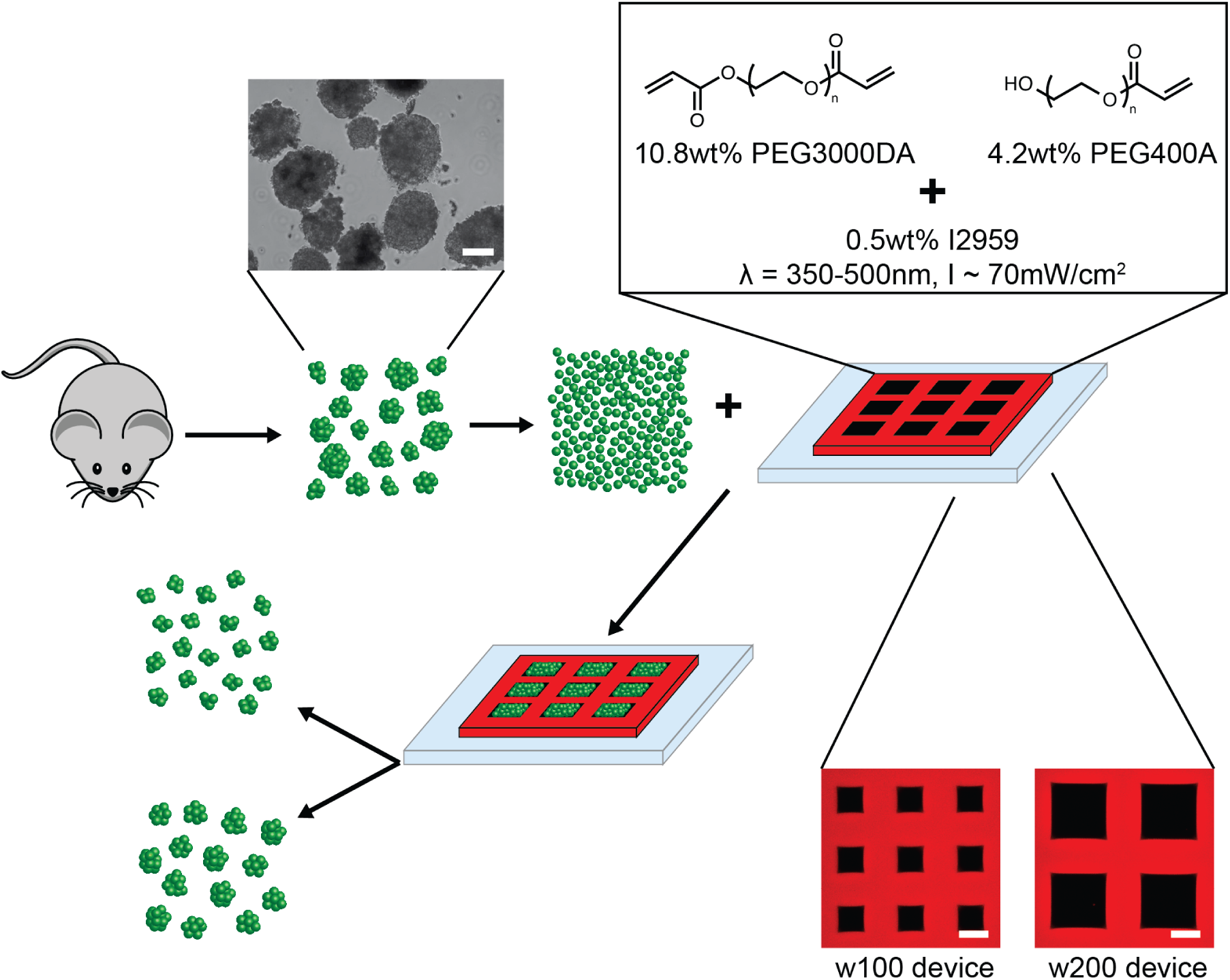
Schematic of primary cell pseudo-islet formation. Murine pancreatic islets are dissociated into a single cell suspension and seeded into poly(ethylene glycol) (PEG) hydrogel microwell arrays with well widths of 100 μm or 200 μm to form pseudo-islets of uniform sizes. Multiwell devices are patterned via a chrome photomask through contact photolithography. PEG-diacrylate and PEG-monoacrylate are photopolymerized with Irgacure 2959 as the photoinitiator with 350-500 nm light between an acrylated glass slide and the photomask. Methacrylated-rhodamine is included in the pre-polymer solution for fluorescent visualization. w100 devices have approximately 1225 wells and w200 devices have approximate 550 wells. Scale bar, 100 μm.

**Figure 2.**
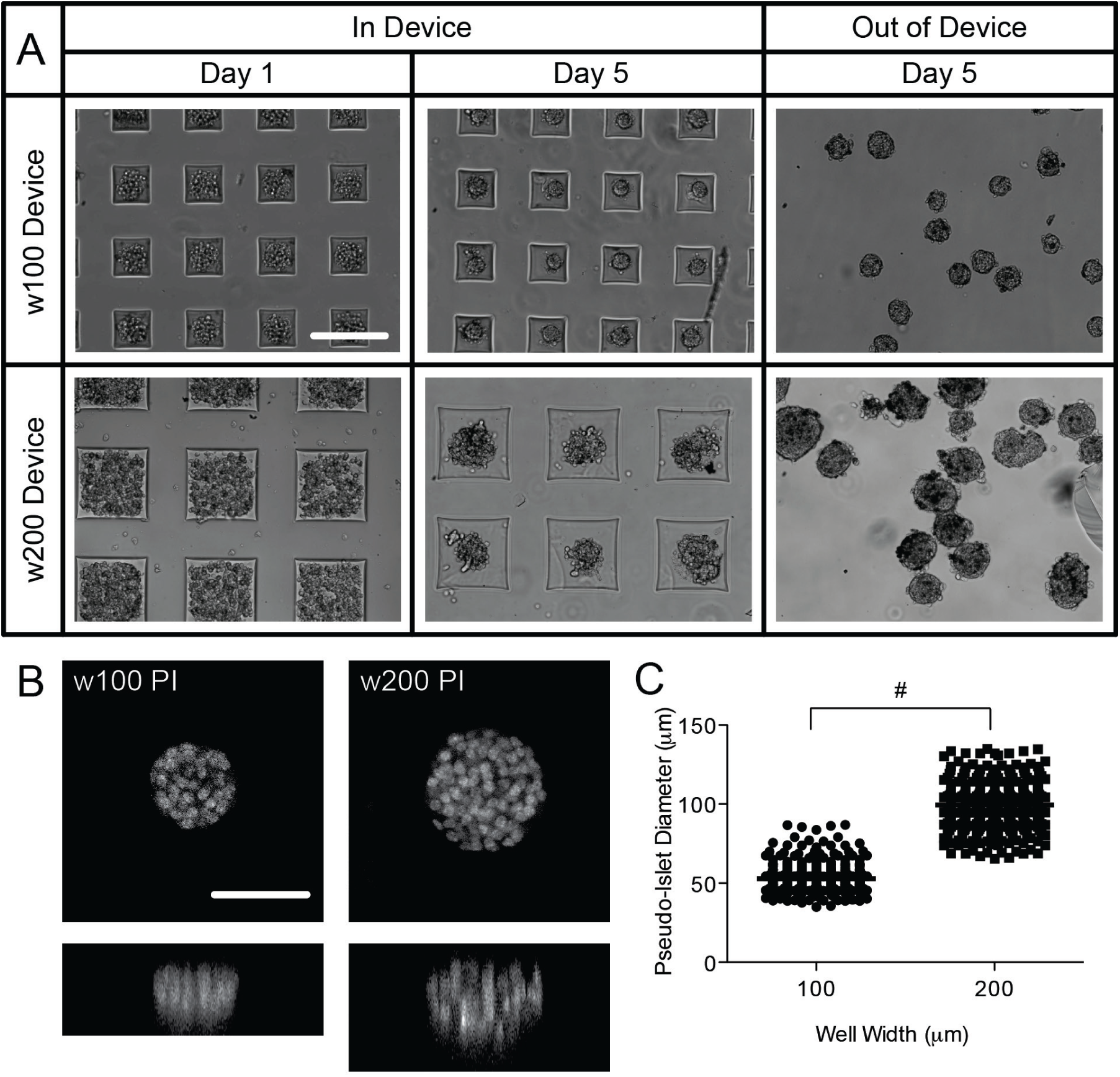
Formation and characterization of pseudo-islets. (A) Brightfield images of primary cells seeded into w100 (100 μm wide) and w200 (200 μm wide) devices immediately after seeding on day 1, aggregated after 5 days of culture in the devices and in suspension after removal from the device. Scale bar, 200 μm. (B) Maximum intensity projections in the XY (top) and XZ (bottom) plane of nuclear staining (DAPI adjusted to greyscale) of pseudo-islets. Scale bar, 50 μm. (C) Diameter of pseudo-islets formed in w100 and w200 devices. n = 328 (w100 PI) or 418 (w200 PI); # p<0.0001.

To confirm that three-dimensional aggregates were created, nuclear staining and image analysis was performed. The pseudo-islets were found to be approximately spherical as evidenced by staining in all three dimensions (x, y, and z) (Figure 2B). Additionally after removal from the devices and image analysis, their diameter was measured to characterize the effect of device dimension on pseudo-islet size. Pseudo-islets from w100 devices (w100 PI) had an average diameter of 50 ± 10 μm while pseudo-islets from 200 μm wells (w200 PI) had an average diameter of 100 ± 15 μm (mean ± standard deviation), and these values were found to be statistically different from each other (p<0.001) (Figure 2C). Over 300 pseudo-islets of each size from 4 different experiments were analyzed.

### 3.2 Cellular composition and connectivity of pseudo-islets

To investigate whether the pseudo-islets had a similar cellular composition compared to the original whole islets, immunofluorescence staining was performed to characterize the three major islet cells types. Similar to whole islets, w100 and w200 pseudo-islets stained positive for insulin, glucagon, and somatostatin indicating the presence of β-cells, α-cells, and δ-cells respectively (Figure 3 and Supplementary Figure 1). As these hormones work together to regulate blood glucose levels, the presence of multiple cell types beyond just β-cells is important. In addition, due to their smaller size, more complete staining throughout the entire cellular aggregate was observed in the pseudo-islets than in whole islets, which display a lack of staining at positions more than approximately 75μm into the islet. The lack of staining in the center of the aggregates is attributed to limitations in diffusion of the antibody through the dense cell mass. The ratio of β-, α-, and δ-cells within the pseudo-islet was evaluated by counting the number of cells that stained positive for insulin, glucagon and somatostatin at 10μm slices throughout the pseudo-islets. Independent of size, ∼70% of cells in the pseudo-islets were β-cells, ∼22-25% were α-cells and the remaining were δ-cells (Supplementary Figure 2). Additionally, no double-labeled cells were observed after examination of single image slices.

**Figure 3.**
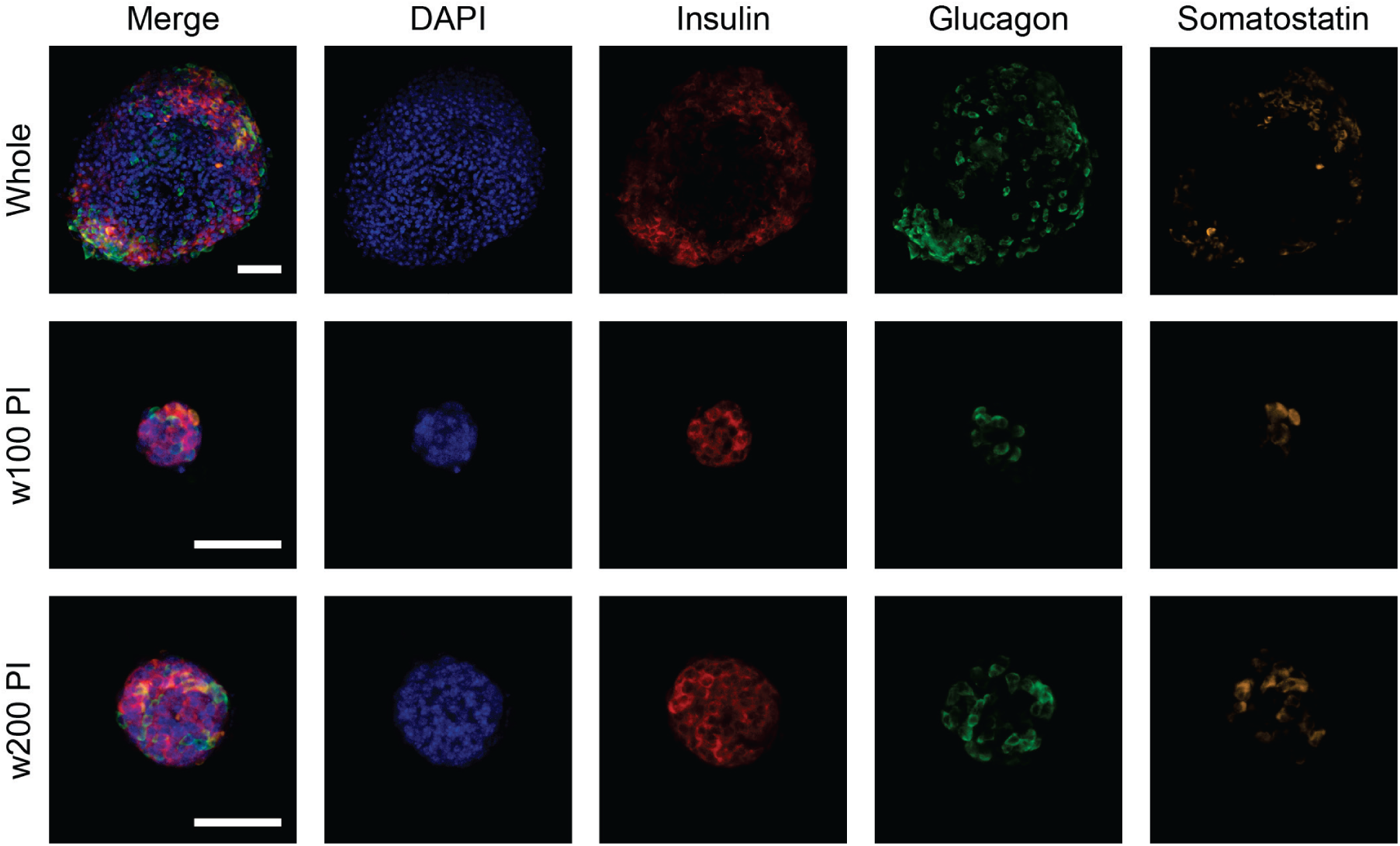
Cellular composition of pseudo-islets. Maximum intensity confocal projections of whole mount immunofluorescence staining of whole and pseudo-islets indicates the presence of β-cells stained for insulin (red), α-cells stained for glucagon (green), δ-cells stained for somatostatin (orange), and cell nuclei labeled with DAPI (blue). Scale bar, 50 μm.

Since cell-cell contact through gap junctions is important for β-cell function, we also completed immunofluorescence staining for connexin-36, found in β-cell-to-β-cell gap junctions. Both sizes of the pseudo-islets stained positive for connexin-36, similar to whole islets (Supplementary Figure 3).

### 3.3 Sustained viability of pseudo-islets over whole islets

As an initial measure of viability, pseudo-islets were stained with a Live/Dead viability kit and imaged with confocal microscopy on days 7 and 14 after islet isolation (Figure 4). Over 14 days in culture, both sizes of pseudo-islets consisted of almost entirely live cells, as indicated by robust calcein staining. At the same time point, whole islets showed signs of increased cell death, as indicated by increased ethidium homodimer staining as well as staining voids, attributed to cells that were dead for greater than 24 hours and lack intact DNA for the dead stain (ethidium homodimer) to bind. Although large portions of the whole islets appear to stain live, this is actually staining on the exterior of the islet, which fills the islet area in the 2-dimensional projections shown in Figure 4. Individual image slices reveal incomplete staining through whole islets which likely results from the inability of calcein and ethidium homodimer to penetrate the inner cell mass combined with metabolism of the labeling molecules before they can diffuse into the islet center (Supplemental Figure 4A). In comparison, the pseudo-islets’ smaller diameter allows for visualization throughout their entire volume.

**Figure 4.**
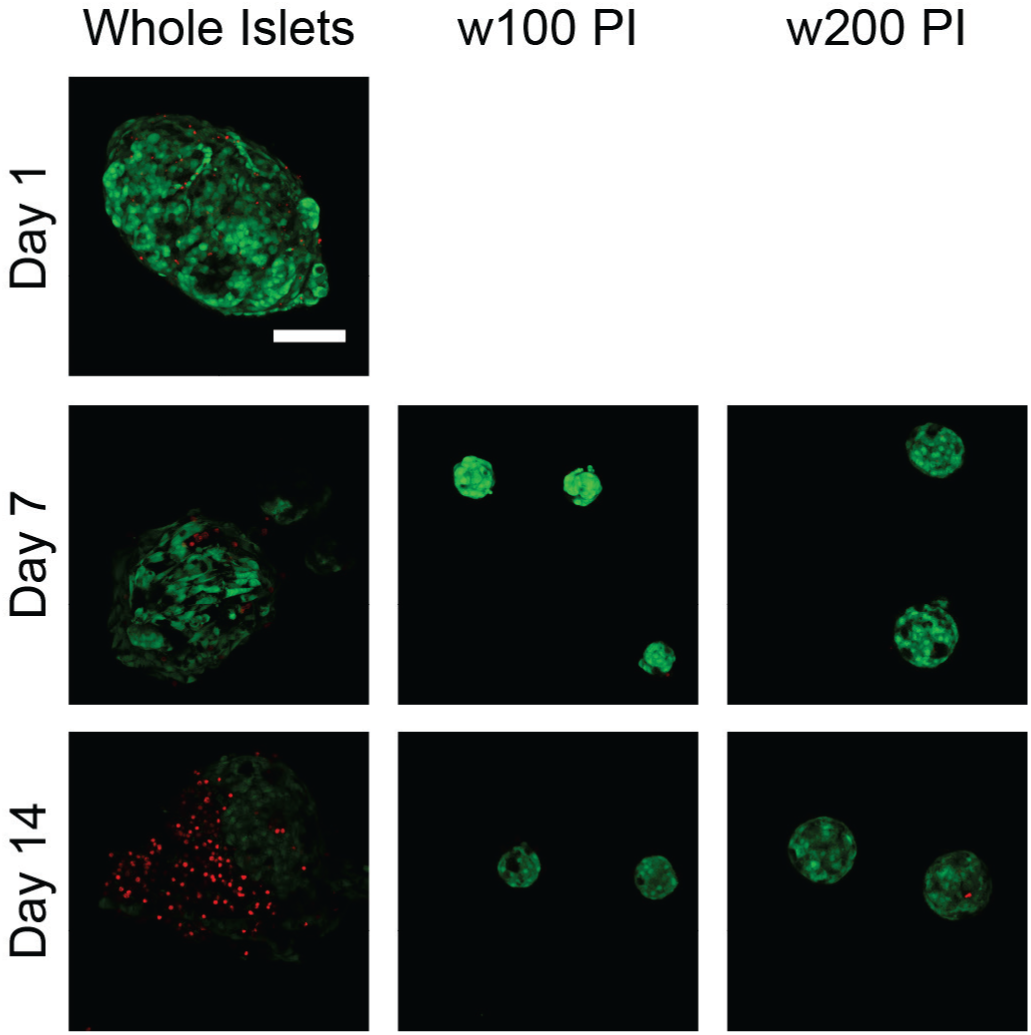
Viability of whole and pseudo-islets via Live/Dead staining. Representative maximum intensity projections from confocal imaging of whole islets one day after isolation and whole and pseudo-islets on days 7 and 14 after isolation. Live cells stain green and dead cells appear red. Scale bar, 100 μm.

### 3.4 Functional behavior of pseudo-islets

The overall functional response of β-cells in pseudo-islets was measured by assaying static insulin secretion in response to glucose challenge over 14 days in culture. To account for the size difference and inherent difference in the number of insulin secreting β-cells, the samples were lysed in order to normalize insulin secretion in the supernatant to the total insulin content of the cell lysate. With extended time in culture, whole islet insulin secretion significantly increased at days seven and fourteen compared to one day after isolation (p< 0.05 after Bonferroni correction) (Figure 5A). This increase in insulin secretion is not due to an improved response to glucose, but rather a decrease in the total insulin content at days 7 and 14 (Figure 5B, p<0.05), which is likely a function of β-cell death in the whole islets. Pseudo-islets of both sizes were found to be responsive to glucose challenge at days seven and fourteen and no statistical difference was found between sizes at either time point. In general, the pseudo-islets exhibit similar behavior to freshly isolated whole islets with respect to their insulin response (not statistically significant). The total insulin content in pseudo-islets is less than that in whole islets since the pseudo-islets are smaller in size. Additionally as expected, the total insulin content in w200 pseudo-islets was higher than w100 pseudo-islets, but both sizes of pseudo-islets had similar amounts of insulin at days seven and fourteen (Figure 5B, not statistically significant).

**Figure 5.**
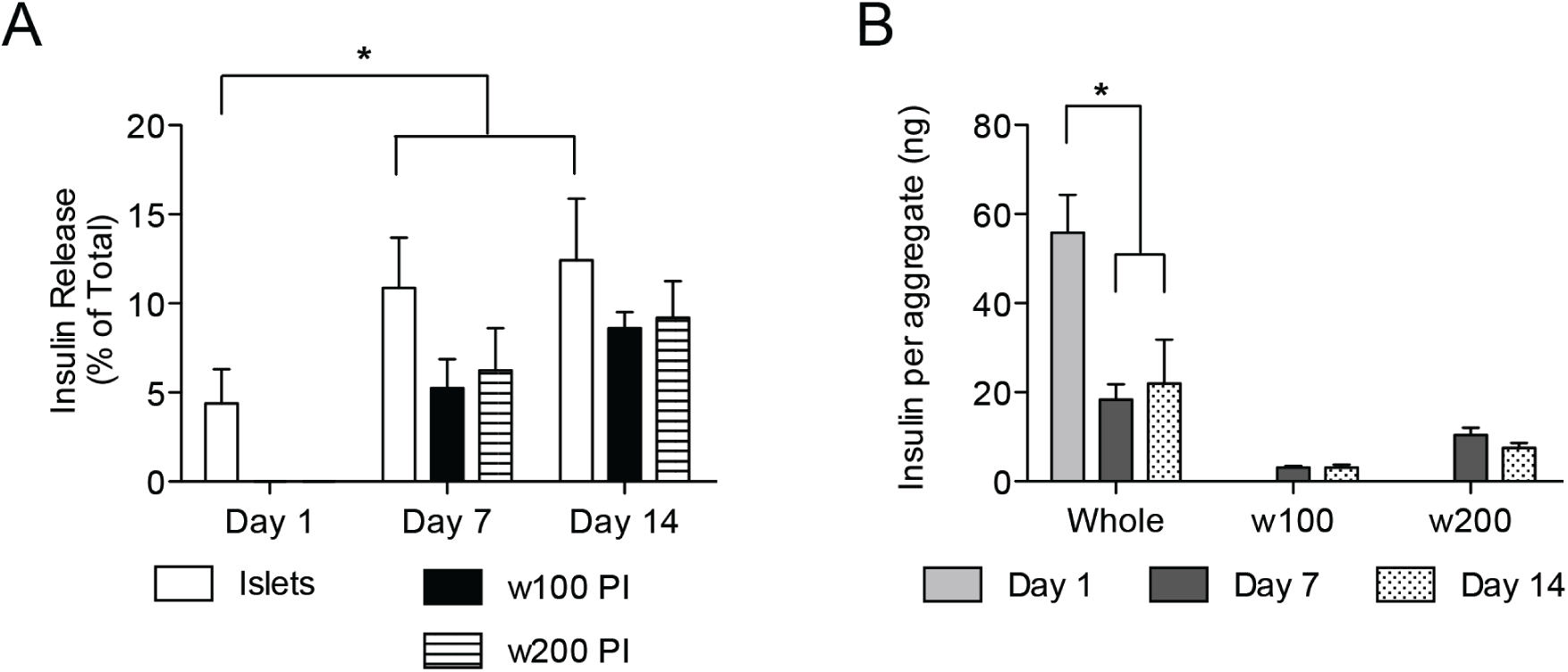
Insulin response from whole and pseudo-islets. (A) Percent of total insulin released in response to 20 mM glucose stimulation from whole islets (white bars), w100 pseudo-islets (black bars) and w200 pseudo-islets (striped bars) over 14 days post-isolation. (B) Insulin content per whole or pseudo-islet over 14 days post-isolation. n≥8; * p<0.05.

### 3.5 Dynamics of pseudo-islet insulin secretion

To further investigate pseudo-islet function, we observed the dynamics of β-cell signaling underlying insulin secretion by monitoring calcium flux with Fluo-4 AM and real-time fluorescence microscopy. The fluorescence intensity traces of a whole islet or individual cells can be examined over time to understand how cells respond in real time to glucose challenge. Here, qualitative observations about islet response were examined by following fluorescence traces over time (Figure 6A). When challenged with 11 mM glucose, healthy cells exhibit regular calcium oscillations. Freshly-isolated islets display regular calcium oscillations one day post harvest (Figure 6A); however with further culture, these oscillations deteriorate into chaotic oscillations by day seven and fourteen (Figure 6A). By comparison, at both of these later time points, w200 pseudo-islets display regular, robust oscillations (Figure 6B).

**Figure 6.**
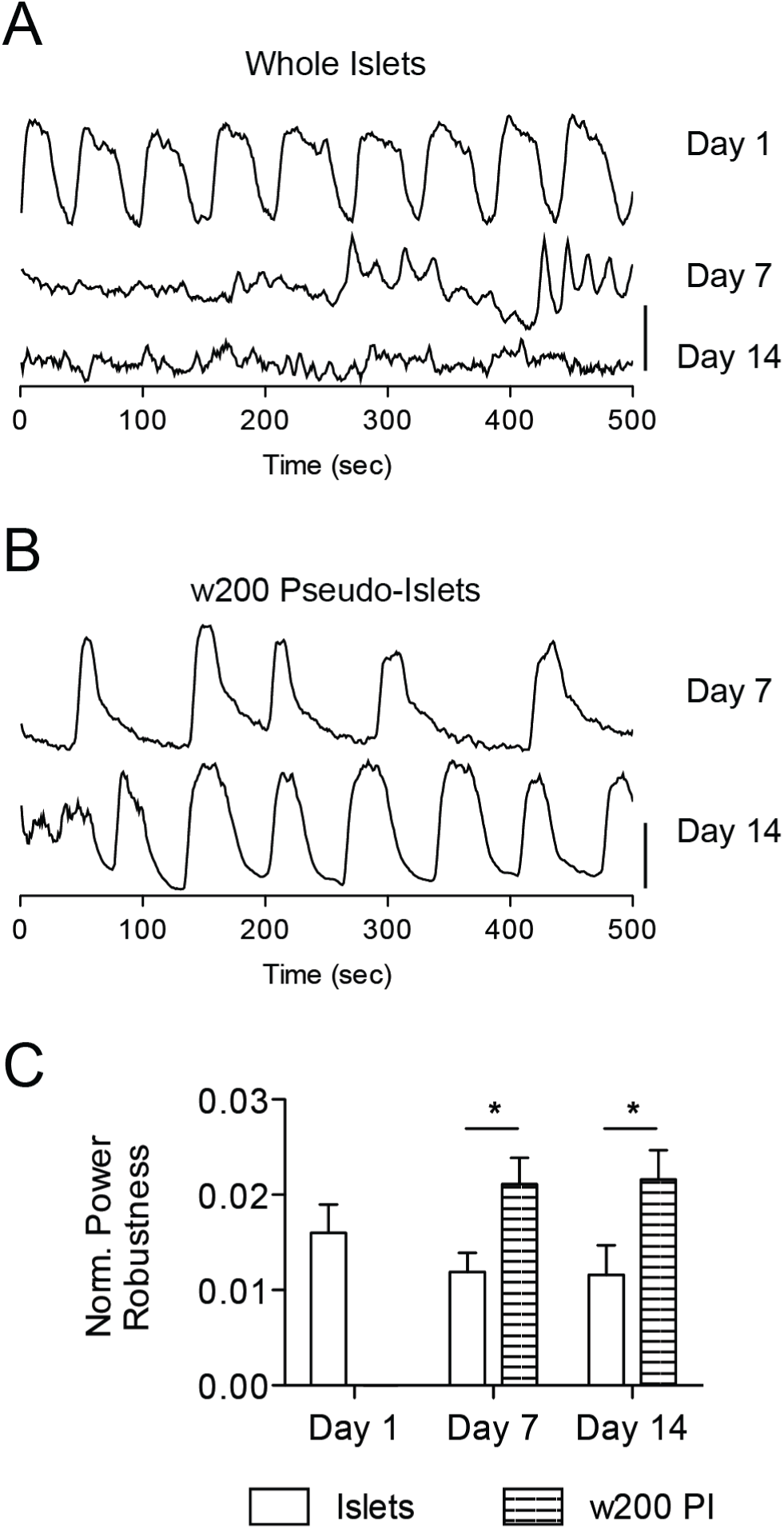
Regularity and robustness of calcium oscillations of whole islets and pseudo-islets at 11 mM glucose. Representative calcium oscillations under high glucose conditions (11 mM) of (A) whole islets at days 1, 7 and 14 and (B) w200 pseudo-islets on days 7 and 14. Time courses are offset for clarity and the vertical line represents a 50% change in fluorescence. (B) Normalized power robustness of calcium oscillations at 11 mM glucose of whole islets (white bars) and w200 pseudo-islets (striped bars) over 14 days post-isolation. n≥8; * p<0.05.

These observations were quantified by calculating a value for the robustness of the period of the oscillations, which is presented in Figure 6C. The decay of regular oscillations in whole islets on day seven and day fourteen is confirmed by a decrease in the normalized power robustness. Similarly, as indicated by the fluorescence traces, the w200 pseudo-islets had statistically higher average normalized power robustness than age-matched whole islets (p<0.05). Pseudo-islet size did not have an effect on the robustness of oscillations, as no difference was observed between the w100 and w200 aggregates (Supplementary Figure 5A).

### 3.6 Coordination of calcium fluctuations within pseudo-islets

Proper cell-cell connectivity is important to maintain a coordinated response of β-cells in an islet under stimulatory conditions and to suppress spontaneous activity under basal conditions. The real-time monitoring of calcium flux with Fluo-4 AM also allowed us to probe the coordination of β-cell response within pseudo-islets. The area of synchronized behavior at high (11 mM) glucose was determined through examining the synchronization of oscillatory calcium flux between cells. Freshly-isolated islets had average synchronization areas that decreased by days seven and fourteen (day 1 vs 14: p<0.05), while both sizes of pseudo-islets at the same time points maintained comparable levels of synchronization, which were similar to the freshly-isolated islets at Day 1 (Figure 7A, Supplementary Figure 5B). Example heat maps of the correlation coefficients across whole islets and w200 pseudo-islets at days one, seven and fourteen are shown in Figure 7B. Warmer tones indicate higher correlation coefficients while cooler tones correspond to lower correlation coefficients.

**Figure 7.**
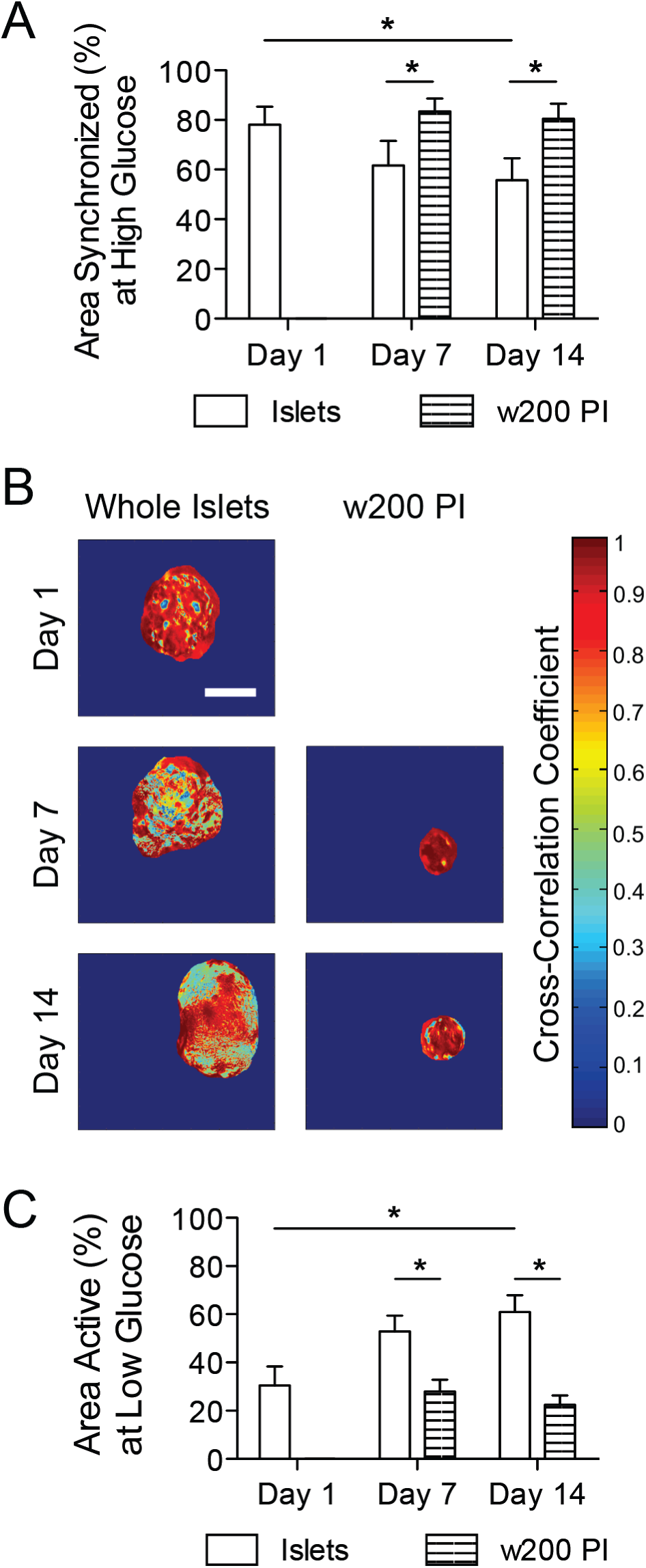
Collective behavior of whole islets and pseudo-islets. (A) Area of whole islets (white bars) and w200 pseudo-islets (striped bars) exhibiting synchronized calcium oscillations at 11 mM glucose over 14 days post-isolation. (B) Heat maps displaying cross-correlation coefficients of whole islets and w200 pseudo-islets on days 1, 7, and 14 post-isolation. Scale bar, 100 μm. (C) Area of whole islets (white bars) and w200 pseudo-islets (striped bars) displaying calcium activity at 2 mM glucose over 14 days post-isolation. n≥8; * p<0.05.

To ascertain the level of activity within a pseudo-islet at low (2 mM) glucose, we calculated the percent active cells within an islet or pseudo-islet. Freshly isolated islets and w200 pseudo-islets at day 14 had low levels of spontaneous activity at low glucose (Figure 7C), indicative of the ability to regulate insulin release. However, higher levels of activity were measured for whole islets on days 7 and 14. The increase in whole islet activity on day 7 and 14, as well as the difference in whole islet and pseudo-islet activity was found to be statistically significant (p<0.05 after Bonferroni correction). Similar to the high glucose measurements, pseudo-islet size did not lead to a difference in the area active (Supplementary Figure 5C), suggesting that both sizes of pseudo-islets were functional in their ability to maintain low levels of activity at low glucose levels.

## 4. Discussion

Due to the absence of a robust cell source for islet transplantation, methods to create β-cell aggregates that display improved viability and function as compared to whole islets are needed to improve the efficiency of islet transplantation. In addition, controlling the size of these β-cell aggregates will be essential as larger islets exhibit lower viability and glucose response [7,8] most likely due to limitations in the diffusion of oxygen and other nutrients into the center of these large cellular clusters [5,43]. Additionally, the recent advances in differentiation of stem cells into β-cells [11–13], renders the ability to effect β-cell coupling through controlled reaggregation favorable in order to generate functional aggregates. Here, we have demonstrated the use of PEG microwell devices to create pseudo-islets of defined sizes from primary murine pancreatic islets. Controlling the size of these pseudo-islets allows for the promotion of β-cell function and viability as compared to whole islets.

The use of microwell arrays has been exploited for use with a range of cell types [44–47], and has been recently utilized to create β-cell clusters [37,38,48]. These hydrogel microwell devices possess many benefits over alternative methods of β-cell aggregation, such as the hanging drop method [33,49] and spontaneous aggregations [14,30,50], including simplicity of fabrication, high fidelity of pattern transfer across a range of sizes and shapes, and the ease of aggregate culture and removal due to use of PEG, a polymer commonly used in biomaterials for it’s high water content and bio-inert nature. We have translated the use of these microwells from mouse insulinoma 6 (MIN6) cells, a β-cell line, to primary islets. Primary islet cells are more limited in number and can require a significant amount of effort and care to isolate clean islets. Additionally, primary cells are more fragile than MIN6 cells and required extensive investigation into culture and dissociation methods to obtain a large number of single, viable cells for seeding into the microwells.

This technique allows us to generate uniform cellular clusters from tissues that are naturally disperse in both size and shape. Here we take whole islets, which range in size from 50 to over 350 μm in diameter and can vary from spherical to ellipsoidal [8,51,52], and produce three-dimensional pseudo-islets whose diameters are tightly controlled and a function of microwell dimensions (Figure 2). The pseudo-islets generated here have average diameters of 50 and 100 μm, which fall below the upper limit of what is generally considered a ‘small’ islet, of 125 μm. Using microwells of a larger size would allow the user to create larger pseudo-islets; however, it is likely undesirable to create larger aggregates and would require a higher number of cells to successfully seed the microwell device, a concern when working with primary cells.

Pancreatic islets contain multiple cells types, which work in concert to control blood glucose levels. Among the three main cell types, β-cells secrete insulin in response to high glucose levels, α-cells secrete glucagon in response to low glucose levels, and δ-cells secrete somatostatin which suppresses both β- and α-cell activity [53,54]. Together these cell types work together in a complex system to control blood glucose levels and altering parts of the multiple pathways in which insulin, glucagon and somatostatin are involved can ultimately affect β-cell function. Specifically, in mice, a functional knockout of the glucagon receptor resulted in changes in glucose metabolism and dysregulation of insulin secretion [55] and deletion of the somatostatin receptor predominately found on β-cells decreased somatostatin’s ability to inhibit insulin secretion [56]. Immunostaining of insulin, glucagon, and somatostatin confirms the presence of these three cell types in our pseudo-islets (Figure 3). In murine islets, β-cells comprise about 75% of the islet mass, α-cells account for about 20% and δ-cells the balance [57,58]. Given the lower levels of α- and δ-cells in murine islets, the immunostaining results confirm that our method does not sort or exclude these cells types from the pseudo-islets. We found that the pseudo-islets have a similar composition to whole islets (Supplemental Figure 2).

In addition to cellular composition, cellular organization may also be important for islet function. Rodent islets have a unique core-mantle organization where β-cells are surrounded by a shell of α- and δ-cells, allowing for high degree of homotypic β-cell contacts. [57,58]. When the whole islet immunostaining images were examined as individual image slices, α- and δ-cells were located on the periphery (data not shown) consistent with these reports. α- and δ-cells that appear to be in the center of the whole islets in Figure 3 are actually on the top surface of the islets, which appears in the center of the islet in the 2-dimensional projections shown. Previous studies have observed single cell reorganization into a core-shell architecture in the spontaneous reaggregation of primary rat islets [31] and in aggregates made from a mixture of cells from a β-cell line and an α-cell line [59]. In our pseudo-islets, α- and δ-cells appeared more often on the aggregate periphery, but a few α- and δ-cells were observed throughout the pseudo-islets. The exact organization of our pseudo-islets is of great interest and necessitates further histological analysis. Additionally, potential changes or improvements in cellular organization as a function of culture time as well as the impact of architecture on pseudo-islet function should be investigated.

One contributing factor in failed islets transplantation is hypothesized to be islet cell death, due to diffusion limitations in tightly packed cellular clusters. Native islets are highly vascularized to assist with oxygen and nutrient transport, but low revascularization is observed when islets are transplanted into the kidney capsule spleen, or liver [4,60,61]. Even whole islets can develop a necrotic core after 24 hours of *in vitro* culture under normoxic conditions [6,62]. We used Live/Dead staining to assess the viability of the pseudo-islets compared to age-matched whole islets. Over fourteen days post-isolation, both w100 and w200 pseudo-islets displayed high levels of viability with very few dead cells (Figure 4). In contrast, whole islets had a large number of dead cells by day fourteen as well as areas with no staining (Figure 4), most likely due to debris from cells that died prior to staining and imaging. While we did not observe a dead core of cells in whole islets, due to the inability to completely stain and image the center of large aggregates as seen in individual image slices from the center of whole islets (Supplemental Figure 4A), the whole islets do display a darker core, indicative of dead or dying cells, when observed under brightfield microscopy at later time points (Supplemental Figure 4B). Due to their smaller size, the calcein and ethidium homodimer signals could be observed throughout the entire pseudo-islet (Supplemental Figure 4A). This high level of pseudo-islet viability was maintained under mild suspension culture conditions, as opposed to perfusion or stirred culture systems designed to assist with molecular diffusion.

The benchmark of β-cell function is insulin secretion in response to elevated glucose levels. Static insulin secretion measurement revealed that pseudo-islets secreted similar levels of insulin as freshly isolated whole islets and retained the ability to secrete an appropriate level of insulin through the manipulations from islets within the pancreas to pseudo-islets (Figure 5A). Additionally, these functional properties did not differ between the two sizes of pseudo-islets studied. In contrast, when whole islets were cultured *in vitro*, the calculated insulin secretion increased relative to total insulin content; however, this was attributed to an overall decrease in the total insulin content (Figure 5B) within the islet because of a loss of viable cells (as observed with the Live/Dead staining). The amount of insulin content in the pseudo-islets did not change between days seven and fourteen, and there was less insulin released from pseudo-islets than whole islets at day 1 due to their smaller size. We hypothesized that our pseudo-islets would have increased viability and function due to their smaller size, and this appears to be validated by our data.

As a complimentary measure to total insulin secretion, a bulk measure of pseudo-islet function, we also studied the dynamics and coordination of the signaling underlying insulin secretion through real-time fluorescence microscopy of calcium flux. Calcium flux into β-cells is initiated by a series of events that begin with glucose metabolism by β-cells, and is required to trigger a burst of insulin secretion from stored insulin granules [22]. Pulsatile insulin is more effective in reducing blood glucose levels than steady insulin secretion [27] and is thus a physiologically relevant measure of β-cell function. Both through qualitative observation and by quantitatively measuring the robustness of the calcium oscillations, the pseudo-islets and freshly isolated islets display regular, robust oscillations, but the whole islets deteriorate into chaotic behavior with increased culture time (Figure 6).

Individual β-cells are heterogeneous in their response to blood glucose levels [16,63], however cell-to-cell coupling through gap junctions allows β-cells to share the electrical signals associated with calcium flux allowing for coordination and synchronization of insulin secretion throughout the entire islet [24,64,65]. While the pseudo-islets must have cell-cell interactions to remain intact after removal from microwell devices, immunostaining also reveals the presence of connexin-36 (Supplemental Figure 3), which comprises β-cell-to-β-cell gap junctions and allows for electrical coupling. At elevated glucose levels, comparison of individual points within the pseudo-islets to the average behavior reveals around 80% of the pseudo-islet area exhibits synchronization similar to freshly isolated islets (Figure 7A). With extended time in culture, the pseudo-islets show higher levels of synchronization than age matched whole islets. At basal glucose levels, β-cell coupling through gap junctions helps to reduce spontaneous calcium flux from more active β-cells such that islets display lower levels of calcium activity [23,29,66]. Freshly isolated islets and pseudo-islets display low levels of activity, as measured through analysis of the standard deviation of β-cell calcium oscillations (Figure 7C). The increase in whole islet activity with increased time in culture reflects the inability of cellular coupling to modulate the behavior of whole islets.

From these measures, pseudo-islets formed in PEG microwell devices have similar functional behaviors to healthy islets and display high levels of synchronization and better suppression of low glucose activity than age-matched whole islets. This occurs independent of pseudo-islet size (Supplemental Figure 5), and due to the smaller pseudo-islet diameter, cell survival is high and in turn β-cell function maintained. If pseudo-islets created through the use of larger microwell devices were tested, we anticipate that their behavior may start to diverge from the w100 and w200 pseudo-islets and more closely resemble the age matched whole islets. While smaller size does lead to improved function, extending the role of size in islet function to single cells is not possible, as the necessity of cell-cell contact in not just coordination, as shown here, but static insulin secretion has been shown [15,17–19], especially the need for 3-dimensional cell-cell contact [42]. Finally, the ability of pseudo-islets to maintain healthy function through two weeks of *in vitro* culture suggests that controlling pseudo-islet size through the use of microwell devices may prove a useful step towards improving islet transplantation.

## 5. Conclusions

In summary, we report on the creation of pseudo-islets from primary murine islets using PEG microwell arrays. Pseudo-islets were made in two reproducible sizes by changing the dimensions of the microwells. The pseudo-islets retained a similar cellular composition as whole islets, staining positive for the hormones secreted from β-, α-, and δ-cells. Over two weeks in *in vitro* culture, pseudo-islet maintained high levels of viability, especially compared to age-matched whole islets. In both static insulin secretion and dynamic calcium measurements, pseudo-islets exhibited behavior similar to freshly isolated islets and whole islet behavior deteriorated over two weeks in culture. Of note, the pseudo-islets tested in this study displayed similar behavior, as both size pseudo-islets are in the range of ‘small’ islets, further demonstrating the importance of size on islet viability and function and demonstrating the benefit of controlled islet cell reaggregation.

## Acknowledgements

The authors would like to thank Philip Pratt at the Diabetes and Endocrinology Research Center (Barbara Davis Center, Aurora, CO) for primary islet isolation. The authors wish to acknowledge Matthew Westacott and Aleena Notary for technical assistance and Nikki Farnsworth for helpful technical discussions. Support for this work was provided by National Institute of Health grants R01 DK076084 (KSA), R00 DK085145 (RKPB); Juvenile Diabetes Research Foundation grant 5-CDA-2014-198-A-N (RKPB); National Science Foundation grant DGE0742434 (THH); and the Howard Hughes Medical Institute (KSA).

## Supplemental Figures

**Supplemental Figure 1.**
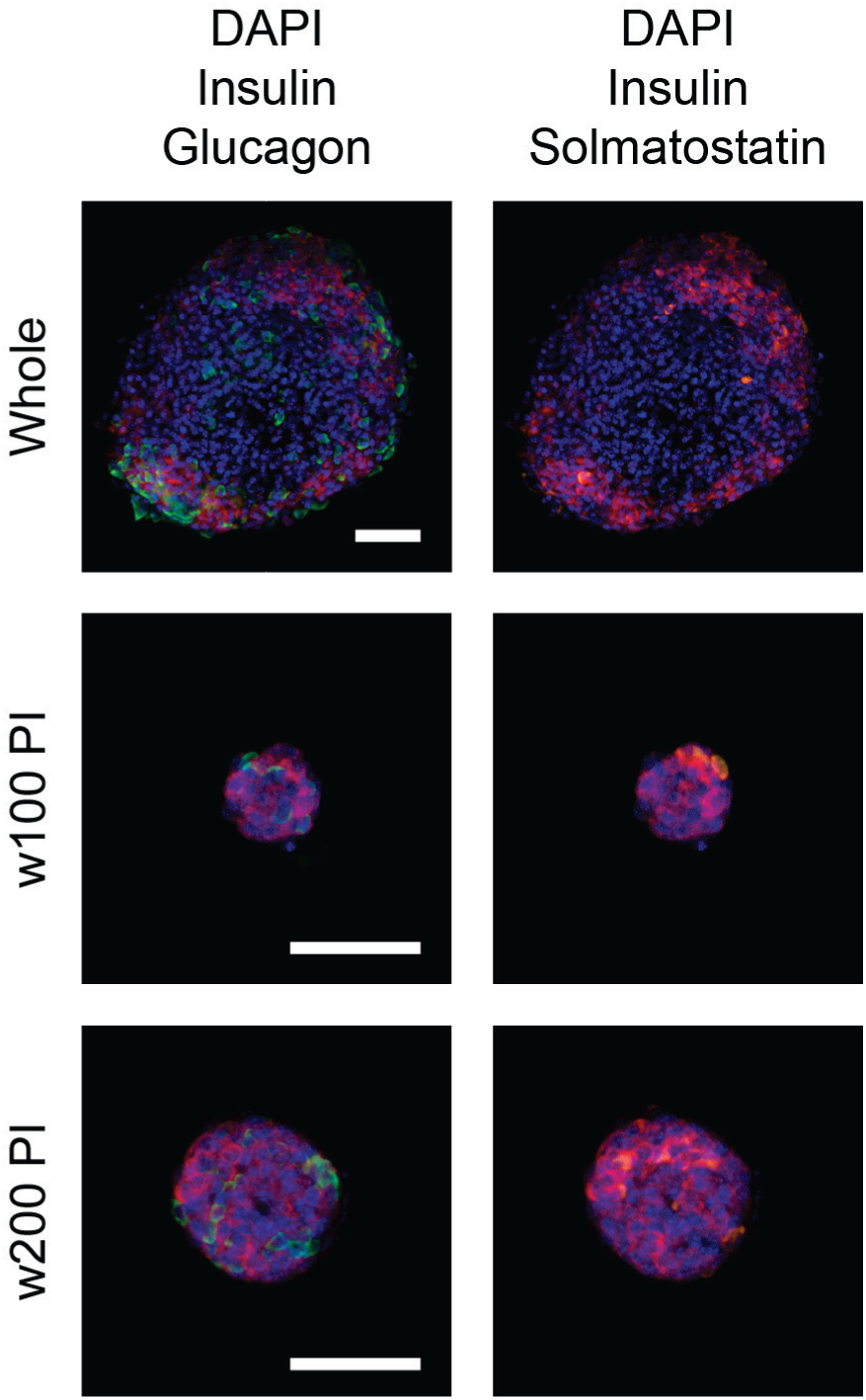
Additional maximum intensity projections of multi-color merged images from whole mount immunofluorescence staining of whole and pseudo islets. β-cells are stained for insulin (red), α-cells stained for glucagon (green), δ-cells stained for somatostatin (orange), and cell nuclei labeled with DAPI (blue). Scale bar, 50 μm.

**Supplemental Figure 2.**
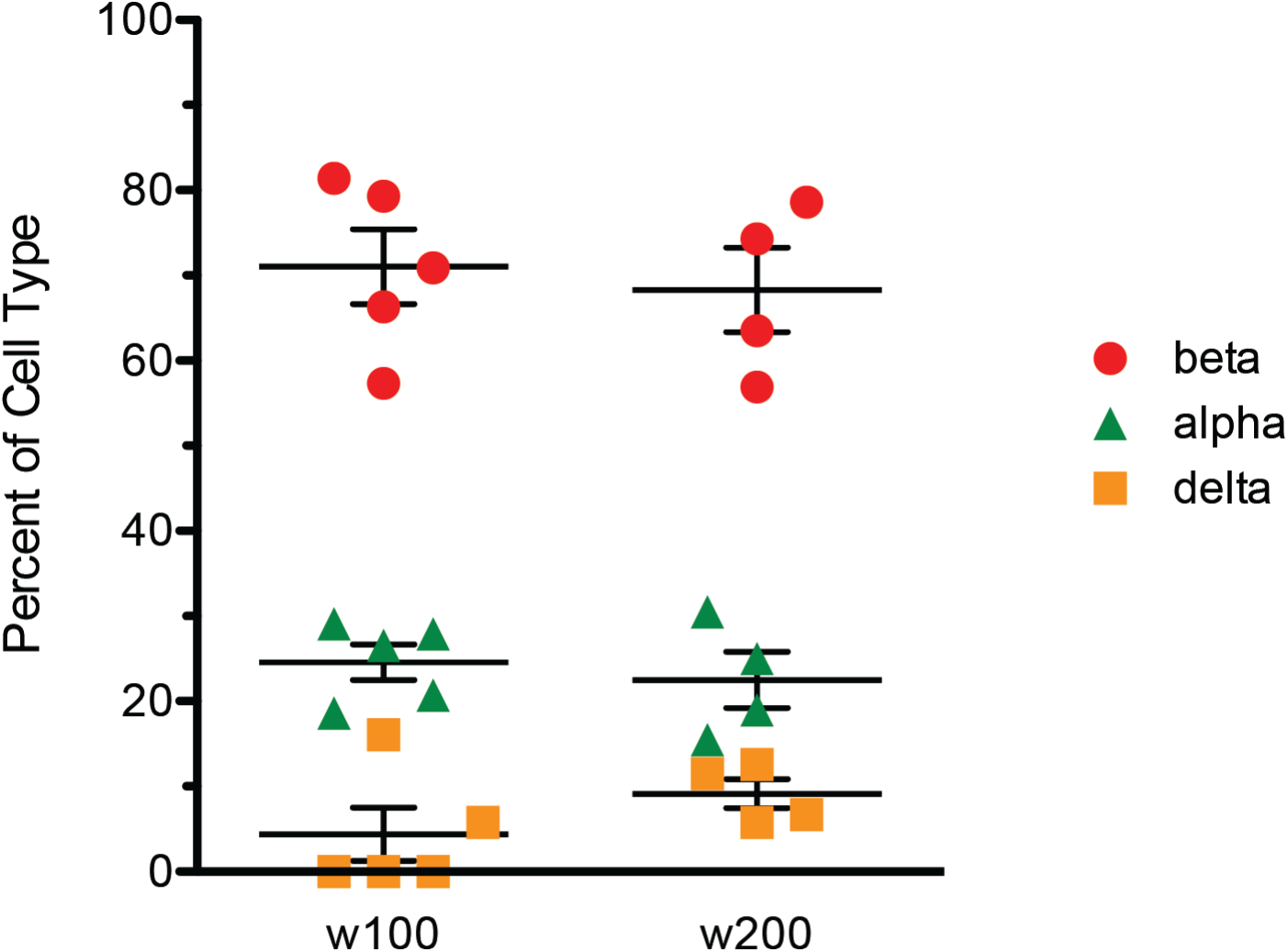
Relative ratios of β-, α-, and δ-cells found within w100 and w200 pseudo-islets as quantified from immunofluorescence images. n=4 (w200 PI) or 5 (w100 PI) and bars represent mean ± standard error.

**Supplemental Figure 3.**
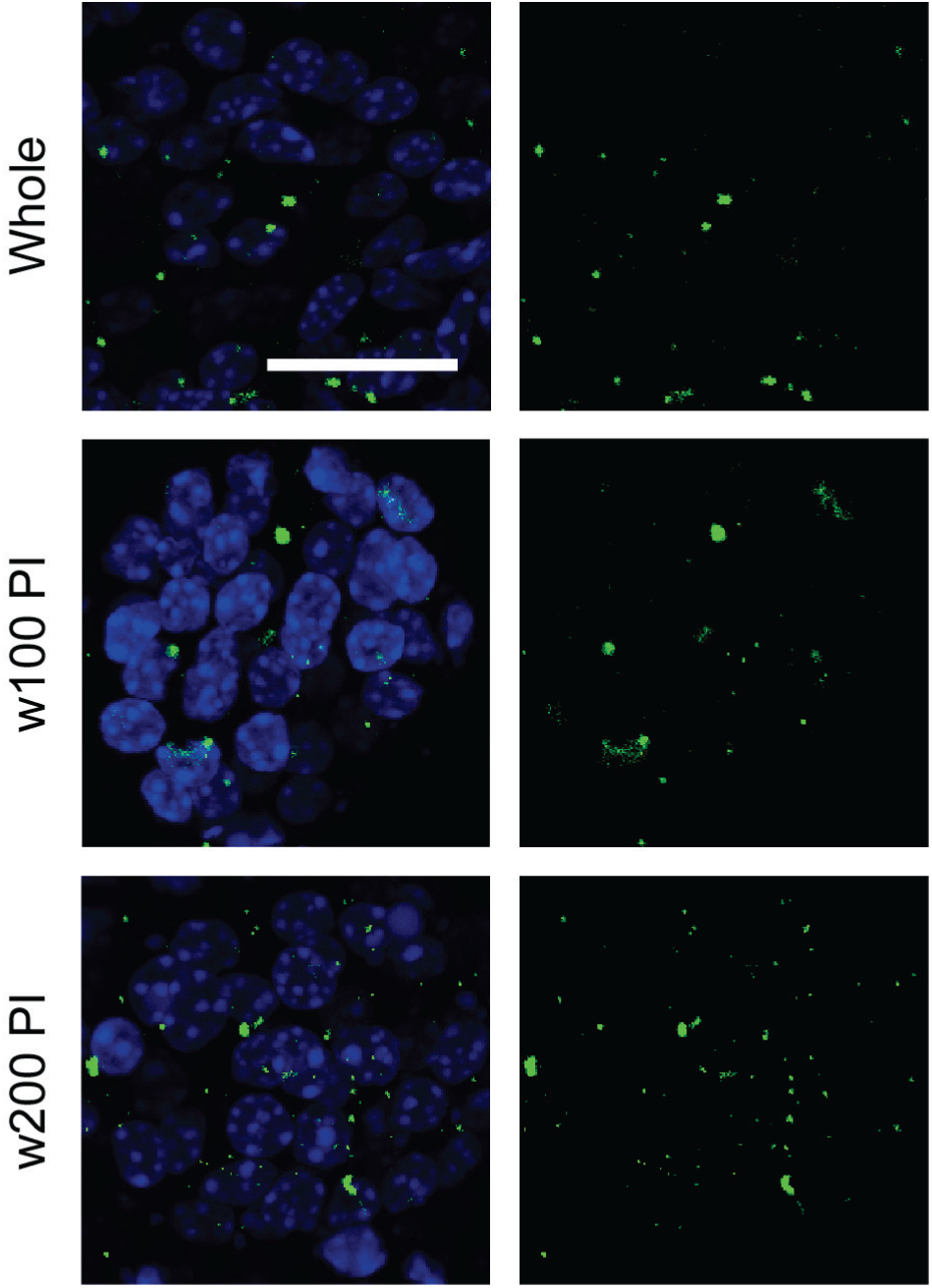
Connexin-36 staining in whole and pseudo-islets. Cx36 stains green and cell nuclei are blue from DAPI staining. Scale bar, 20 μm.

**Supplemental Figure 4.**
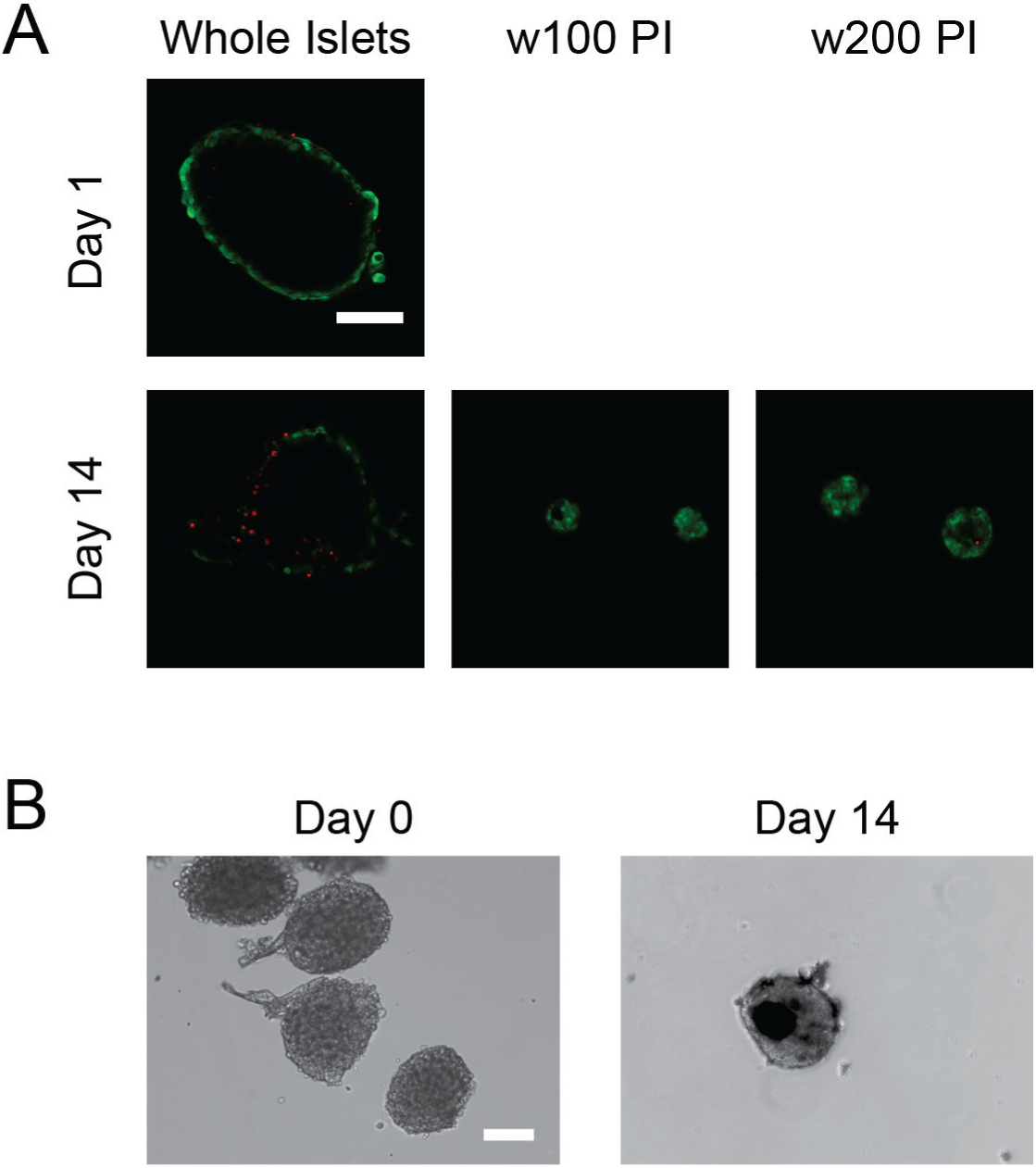
Whole and pseudo-islet viability observed with fluorescent Live/Dead staining and brightfield microscopy. (A) Example confocal z-slices from Live/Dead viability staining of whole islets at day 1 and whole and pseudo-islets at day 14 post-isolation. Whole islets display a lack of staining at the center at both time points, while pseudo-islets can be stained and imaged through the aggregate. (B) Freshly isolated islets (day 0) display consistent translucency. 14 days later, whole islets display a dark core, most likely dead cells. Scale bar, 100 μm.

**Supplemental Figure 5.**
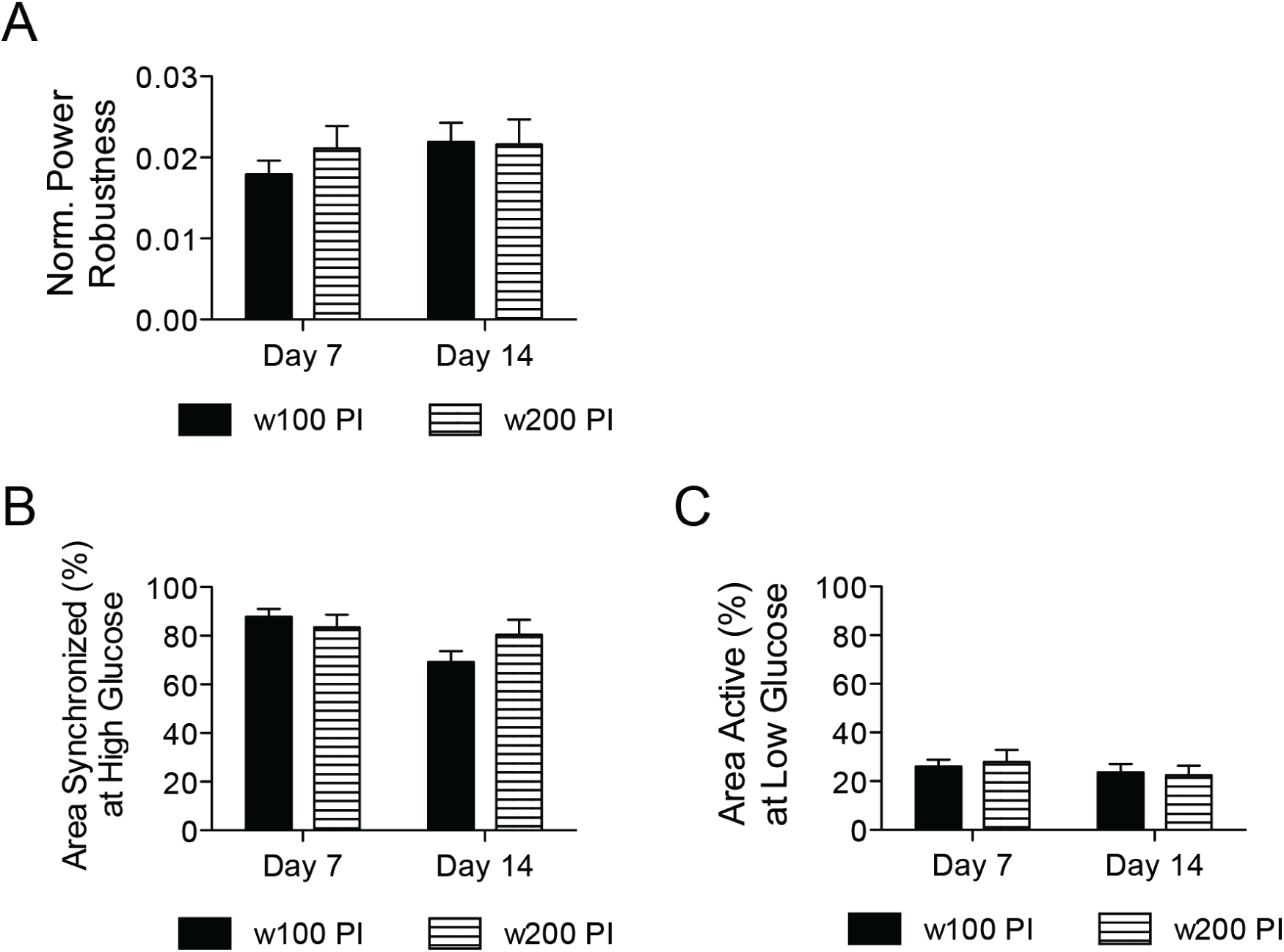
Pseudo-islet size does not affect oscillatory calcium response. (A) Normalized power robustness at 11 mM glucose, (B) area of pseudo-islets exhibiting synchronized calcium oscillations at 11 mM glucose and (C) area of pseudo-islets displaying calcium activity at 2 mM glucose of w100 (black bars) and w200 (striped bars) pseudo-islets at days 7 and 14 post-isolation. n≥8.

